# Neural mechanisms and effects of acute stress on athletes’ unfair decision-making behavior

**DOI:** 10.1101/2025.07.25.666737

**Authors:** Jiayuan Ji, Huiling Wang, Shiyu Wang, Yutong Ye, Yitong Zhang, Lin Li

**Affiliations:** Key Laboratory of Adolescent Health Evaluation and Exercise Intervention, Ministry of Education, East China Normal University, Shanghai 200241, China; School of Physical Education and Health, East China Normal University, Shanghai 200241, China; Center of P.E., Xi’an Jiaotong University, Xi’an, 710049, China

**Keywords:** acute stress, athletes, sense of injustice decision-making, neural mechanisms, functional near infrared spectroscopy (fNIRS)

## Abstract

Acute stress may disrupt decision - making by affecting cognitive and emotional processing. The behavioral and neural mechanisms of this in athletes are unclear. This study explored how acute stress impacts athletes’ unfairness - related decision - making and its neural basis.

Forty participants (20 university athletes and 20 non - athlete students) were randomly assigned to a stress group or a control group. Using functional near - infrared spectroscopy (fNIRS), the study monitored the prefrontal cortex (PFC) and temporoparietal junction (TPJ) blood oxygenation during an ultimatum game task after inducing acute stress via the Maastricht Acute Stress Test (MAST).

Athletes under stress were more accepting of relatively unfair decisions than non - athletes. This was linked to lower activation in the frontal - eye areas (CH15), supramarginal gyrus (CH38), and somatosensory association cortex (CH67), and higher activation in the primary motor cortex (CH64) in athletes. The increase in acceptance efficiency correlated significantly with the reduced CH38 activation (Rho = - 0.425) and increased CH64 activation (Rho = 0.499).

Long - term exercise likely enhances PFC - TPJ functional integration, helping athletes adopt adaptive strategies under acute stress. These findings offer insights for developing stress management and neuromodulation training programs for athletes.

## Introduction

A sense of fairness is one of the key topics in human behavior research. When individuals perceive that fairness principles are violated, it may trigger irrational behaviors.^[1, 2]^ or negative response behaviors^[3]^. Decision making based on subjective fairness perception is known as unfairness perception decision making and is commonly measured by Ultimatum Game (UG) and Dictator Game (DG). In competitive sports, athletes often encounter various types of acute stress events such as individual performance errors, sudden changes in climate, or accidental injuries. If athletes judge the fairness of referee’s decision and opponent’s provocation due to acute stress, they may have irrational reaction which may affect their individual performance or even team performance. Therefore, exploring athletes’ perception of unfairness and decision-making under acute stress can help optimize their on-field coping strategies and reduce the risk of stress-induced decision-making, which is of great value to the practice of competitive sports.

Existing research has found that acute stress can have an impact on sense-of-unfairness decisions. For example, Cano-López et al. (2016) found that individuals under conditions of acute stress are more inclined to reject unfair programs (Cano-López I et al., 2016), hypothesizing that this phenomenon is related to acute stress-induced emotional fluctuations (e.g., anger or anxiety), which enhances an individual’s sensitivity to unfairness^[4, 5]^. However, there are also studies that have reached the opposite conclusion. For example, Takahashi et al. (2007) found through the use of a dictator game task (DG) that individuals accept unfair programs more generously after acute stress^[6]^. Research on athlete populations further revealed the complex effects of acute stress on decision-making patterns: college athletes specializing in volleyball showed a decrease in the correctness of cognitive decision-making but an increase in the correctness of intuitive decision-making after acute stress^[7]^; high-level athletes had higher decision-making accuracy at high arousal levels, whereas low-level athletes performed better at low arousal levels^[8]^. However, there is still a lack of empirical evidence for the stress-influenced effects of decision-making specifically targeting athletes’ sense of unfairness, and its neural mechanisms need to be further explored.

Recently, functional brain studies have provided a neural basis for understanding the relationship between stress and decision making. The Default Mode Network (DMN) and the Central Executive Network (CEN), among others, have been shown to be involved in the acute stress response^[9–12]^. It mainly involves the medial prefrontal cortex (mPFC), the inferior parietal lobule (IPL), the dorsolateral prefrontal cortex (dlPFC), the dorsomedial prefrontal cortex (dmPFC) and the frontal cortex (FEF)^[13]^. In addition, it has also been found that acute stress primarily activates the prefrontal cortex^[14, 15]^ and the temporoparietal joint area^[16]^. Researchers have found that acute stress interferes with the function of the prefrontal cortex by activating the Hypothalamic-Pituitary-Adrenal Axis (HPA axis) and the sympathetic nervous system^[17, 18]^. The function of prefrontal cortex is interfered with, resulting in decreased inhibitory control and emotion regulation^[14, 18]^, which in turn affects fairness judgment^[19, 20]^. For example, a Transcranial Magnetic Stimulation (TMS) experiment showed that participants were more inclined to accept unfair allocation schemes when the right dorsolateral prefrontal lobe was suppressed^[21]^, suggesting that changes in the dorsolateral prefrontal lobe function may be a key factor in the decision-making process affected by acute stress. stress affecting decision-making. Currently, the effects of prolonged exercise on cognition, mood and brain function have received a lot of attention from many scholars, and it has been found that prolonged exercise can enhance the functional connectivity of the prefrontal-striatal circuits, which can improve cognitive control and emotion regulation^[22]^. Meanwhile, long-term exercise training was also found to effectively attenuate stress-induced amygdala and hippocampal neurological damage, thereby accelerating the neurological recovery process^[23, 24]^, suggesting that the exercise experience may remodel neuromodulatory functions. Whether such changes in neuroplasticity induced by exercise experience may mitigate the effects of stress on decision-making remains unknown.

In order to reveal the athletes’ sense of unfairness decision-making and their performance after acute stress, and to investigate its neural mechanisms, the present study used near-infrared spectroscopic imaging (fNIRS) to monitor the blood oxygen signals in the key regions of the PFC and the TPJ of the athletes and ordinary college students when they completed the UG task after acute stress, and aimed to test the following hypotheses by a randomized controlled experimental design: acute stress may affect the athletes’ sense of unfairness decision-making and would trigger changes in the activation patterns of the PFC and TPJ in the brain. This study is intended to provide a scientific basis for the optimization of athletes’ psychological regulation and stress management strategies, to enhance athletes’ decision-making efficacy and competitive performance on the field of play, and to enrich the theoretical model of stress-related decision-making and expand the perspective of neuroscientific research in sports.

## Methods and Materials

### Participants

To estimate the required sample size for this study, we conducted a priori analysis using the G*Power 3.1 software^[25]^. Since the research design was a 4 (groups: athlete stress group; athlete sedentary group; general college student stress group; general college student sedentary group) x 2 (time: pre-test; post-test) experimental design, the test family chosen was F-test-analysis of variance (ANOVA), with the medium effect size (f) set at 0.30 and the statistical validity (1 - β) at 0.80, which means that there is an 80% probability of correctly rejecting a test. 80% probability of correctly rejecting a false null hypothesis. Meanwhile, the significance level (α err prob) was set at 0.05, which indicates that the study is willing to accept a 5% risk of incorrectly rejecting the true null hypothesis. Based on these parameters, the total sample size (N) was calculated to be 36 for this experiment.

In this experiment, 40 college students were randomly recruited from a university in Shanghai, including 20 college athletes and 20 ordinary college students with no sports training experience, half of whom were men and half of women, and were randomly assigned to the stress group and the sedentary group to form four experimental groups, namely, the athletes’ stress group, the athletes’ sedentary group, the ordinary college student’s stress group, the ordinary college student’s sedentary group, and 10 people in each group, with a balanced distribution of genders, and with an age between 17 to 24 years old.

Recruited athletes are required to have at least 5 years of professional and systematic sports training experience. The general college student group, on the other hand, had no long-term professional and systematic sports training experience. All participants were right-handed, had normal visual acuity or corrected vision, no color blindness or color deficiency, and no history of mental or physical illness. The study followed the ethical guidelines of the Declaration of Helsinki and was approved by the Human participants Committee of East China Normal University (Approval No. HR2-0125-2025), and participants signed an informed consent form prior to the experiment and were paid accordingly at the end.

### Experimental Procedure

Upon arriving at the laboratory, participants first read the informed consent document and complete the relevant questionnaires. The experimenter then provides a detailed overview of the experimental procedures, ensuring that each participant has a thorough understanding of the experiment. Following this, the experiment officially commences. Participants sat in front of the computer and put on the NIR electrode caps, heart rate bands, and blood pressure monitors. Subsequently, a 3-minute resting state was entered, and the main subject collected the heart rate and blood pressure at rest. After the resting state, participants began to complete the ultimatum gaming task. At the end of the task, the stress group performed the Maastricht Acute Stress Test, which induces stress through ice water immersion and a mental arithmetic task combined with camera gaze to enhance the stress effect, and the sedentary group performed a 4-minute sedentary break. Blood pressure and heart rate data were continuously collected from the participants throughout the task. Finally, the participants sat in front of the computer again to complete the second ultimatum gaming task. The specific experimental flow is shown in Figure 1.

**Fig. 1.**
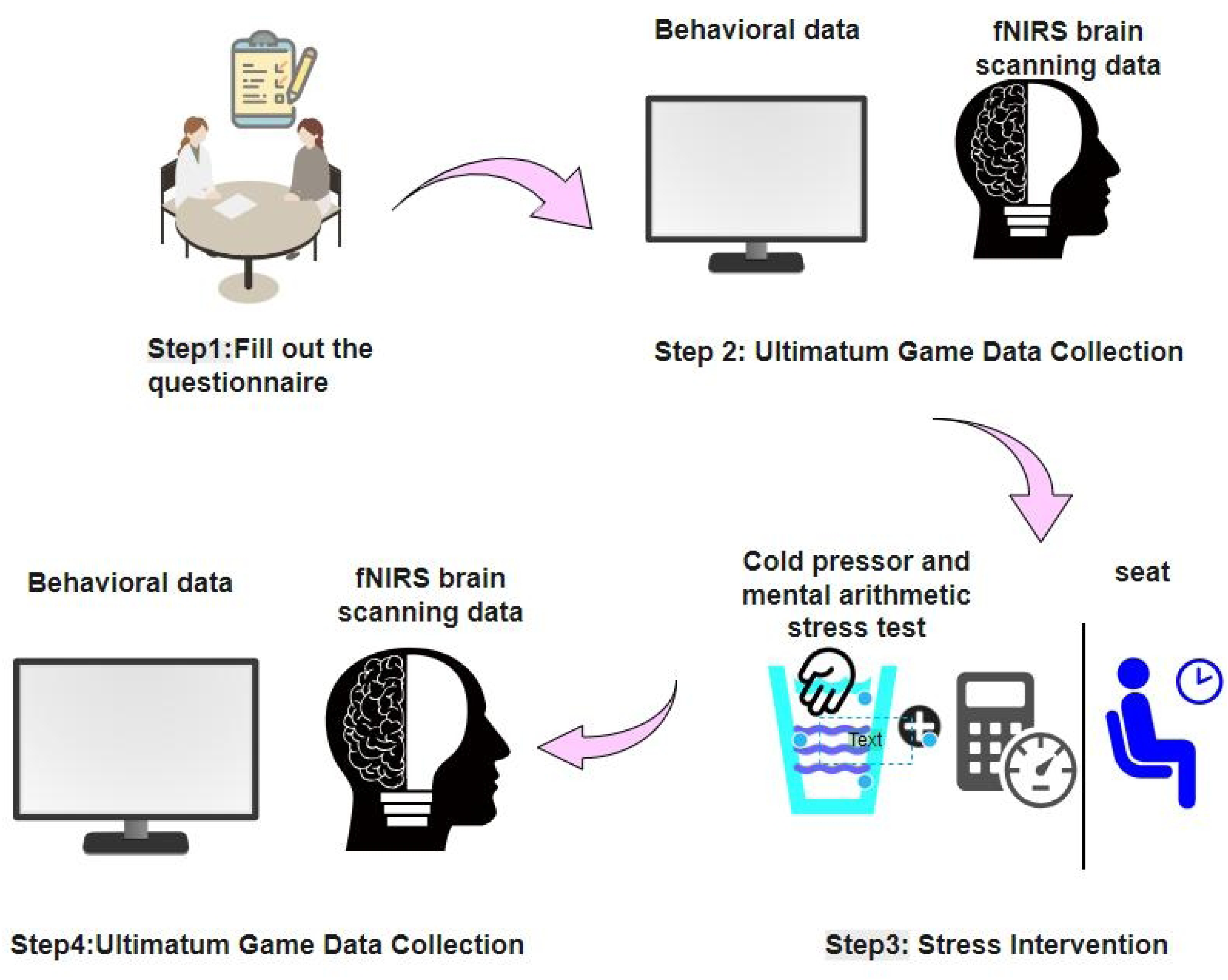
Flowchart of the experiment.

### The Maastricht Acute Stress Test (MAST)

To induce acute stress, this study used a modified version of the Maastricht Acute Stress Test^[26, 27]^, which was designed to alternate between an ice-water immersion and a mental arithmetic task lasting a total of 4 minutes. First, participants were required to complete the ice water task, i.e., to immerse their left hand in cold water of about 0 to 4 degrees Celsius for 1 minute. Then the hand was removed from the water, placed on a towel on the table, and the mental arithmetic task (e.g., subtracting 17 from 2043) was performed, again for 1 minute. The cold water task+ The mental math task consists of one round, and two rounds are performed. Throughout the process, a video camera was pointed at the participants, informing them that their facial expressions would be recorded throughout.

### The Ultimatum Game

This experiment used an ultimatum game task (Ultimatum Game, UG,^[28]^) prepared by EPrime 2.0. In the task, the experimentally proposed observed participants acted as responders and made decisions interactively with a virtual proposer via a computer monitor. There were three allocation schemes (extremely unfair: 1:29-5:25; relatively unfair: 10:20-14:16; and absolutely fair: 15:15), and each type of scheme was pseudo-randomized to be presented 10 times each (total number of trials = 30). The ecological validity of the experiment was strengthened by completing 3 practice trials before the formal experiment and by informing the virtual incentive rule: accepting the proposal resulted in a reward of the corresponding amount, while rejecting it resulted in zero gain for both parties (the actual payoff was irrelevant to the decision).

The flow of the experiment is shown in Figure 2: first, the “+ “ gaze point appears on the screen for 2 seconds. Then the screen displays the allocation proposal, which lasts for 4 seconds. Next, the subject needs to make a decision by pressing the key (F/J) to go for accepting or rejecting the proposal, which lasts for 10 seconds. After that, the results of the choice were displayed on the screen for 4 seconds, including the subject’s income and the peer’s choice. This was followed by a mood assessment phase, in which participants were asked to rate their current mood (“very sad” = 1 to “very happy” = 9) at the push of a button within 4 seconds. Finally, the “+ “ gaze point screen appeared again, signaling the start of the next round. Each round takes about 25 seconds to present and lasts about 13 minutes in total.

**Figure 2.**
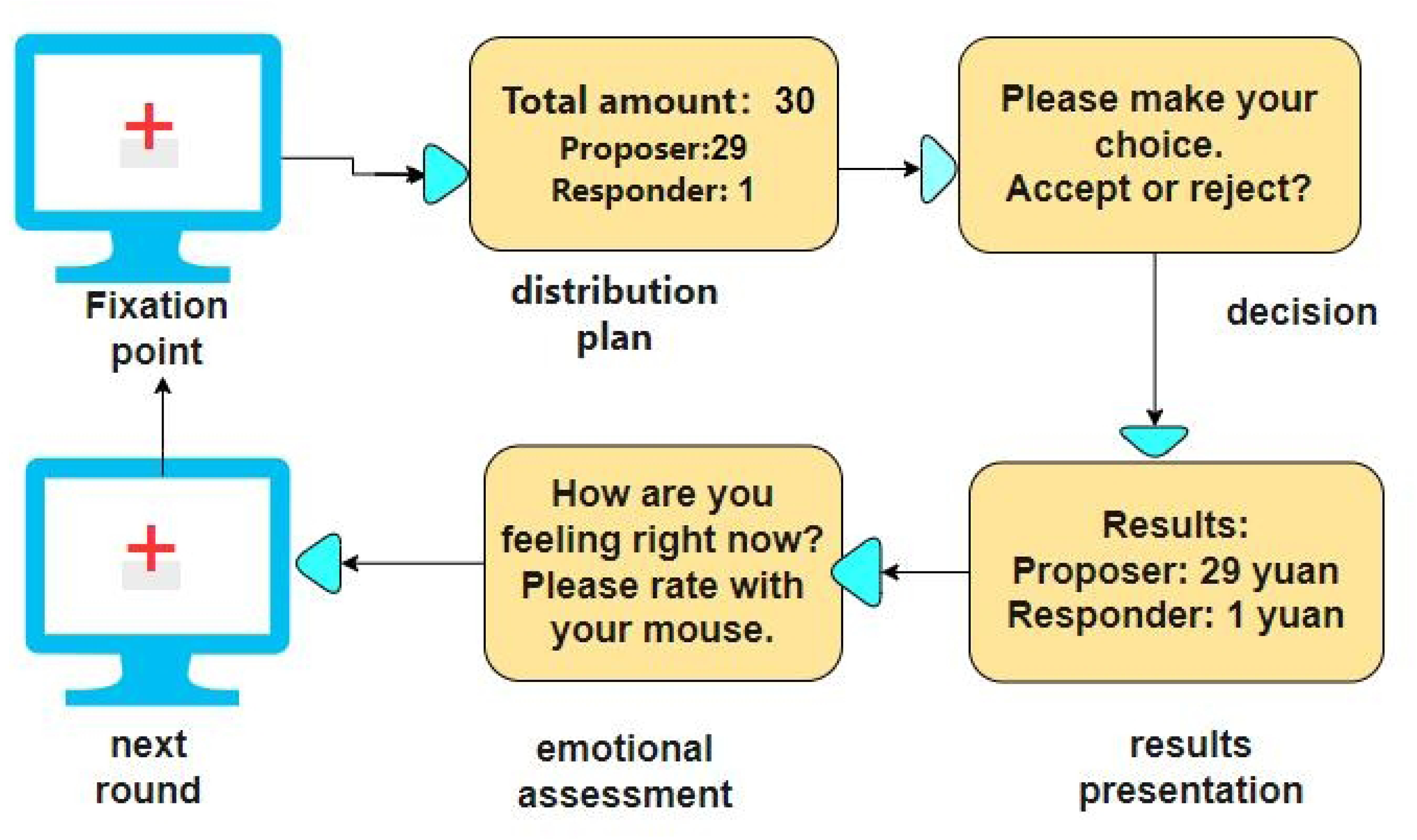
Task flow chart.

### Measurement questionnaire

According to previous studies, personal sense of unfairness is susceptible to many social factors such as gender, personality, and emotion^[29–31]^. Therefore, in this study, the questionnaire method was used to investigate the above indicators and control for irrelevant factors.

### Fairness Perception Questionnaire (Colquitt’s Organizational Justice Scale, OJS)

Based on Colquitt et al. (2001) revised^[32]^ containing four dimensions of procedural fairness (7 questions), distributive fairness (4 questions), interpersonal fairness (4 questions), and informational fairness (5 questions) on a 5-point scale (1=never, 5=frequent), with higher total scores indicating fewer experiences of unfairness.

### Short Version Personality Scale (Chinese Five Personality Scale 2018, CFPS2018)

The short version of the personality scale CFPS2018^[33]^, has 5 dimensions, and its design is based on the Five - Factor Model (FFM), which measures the personality trait scores of dutifulness, extroversion, affinity, openness, and emotional instability, respectively. The questionnaire consists of 15 questions, with 1-5 raw options, and the total dimension score reflects the strength of the trait, with higher scores indicating higher performance.

### General Sense of Power Questionnaire (SPS)

Developed by psychologists Anderson, John and Keltner 2012^[34]^ to assess an individual’s sense of power in social interactions. Participants were asked to score each question based on their true feelings. The scores ranged from 1 (very non-conforming) to 7 (very conforming), with higher scores indicating a greater sense of perceived power for that individual.

### Risk Attitude Scale (RAS)

Developed by Weber (2002)^[35]^ to assess an individual’s attitudes toward risk across multiple domains. The scale aims to understand individuals’ risk acceptance and decision-making styles in different contexts and covers six main domains, each of which reflects an individual’s attitudes and tendencies towards relevant risks through a series of questions.

### Depression Anxiety Stress Scales - 21 (DASS21)

The Depression Anxiety Stress Scale (DASS) is a widely used psychological assessment tool for measuring an individual’s level of depression, anxiety and stress^[36]^. Developed by Lovibond et al. in 1995, the original version was the DASS42, and at a later stage, in order to simplify it and improve its usefulness, the DASS21 was produced, which contains 21 entries divided into three subscales: depression, anxiety and stress, each of which consists of 7 entries and is scored on a 4-point scale.

### fNIRS acquisition

In this study, functional near-infrared spectroscopy (fNIRS) data were continuously monitored using a Hitachi ETG-7100 near-infrared spectroscopic imaging system (Hitachi Medical Corporation, Japan). The system operated at wavelengths of 695 nm and 830 nm, with a sampling frequency of 10 Hz, and a probe array covering a 3 × 5-channel layout (70 channels in total) centered on Fpz (International 10-20 System), with a distance of 3 cm between probes. The device recorded signals of changes in blood oxygenation in the brain during the task. The study used a 3D digitizer to determine the locations of the probes and channels, and identified the corresponding Brodmann areas as regions of interest, including the prefrontal, left and right temporoparietal joint areas (Speer and Boksem, 2019). The BrainNet Viewer^[37]^ was used for brain visualization in this study (Figure 3).

**Figure 3.**
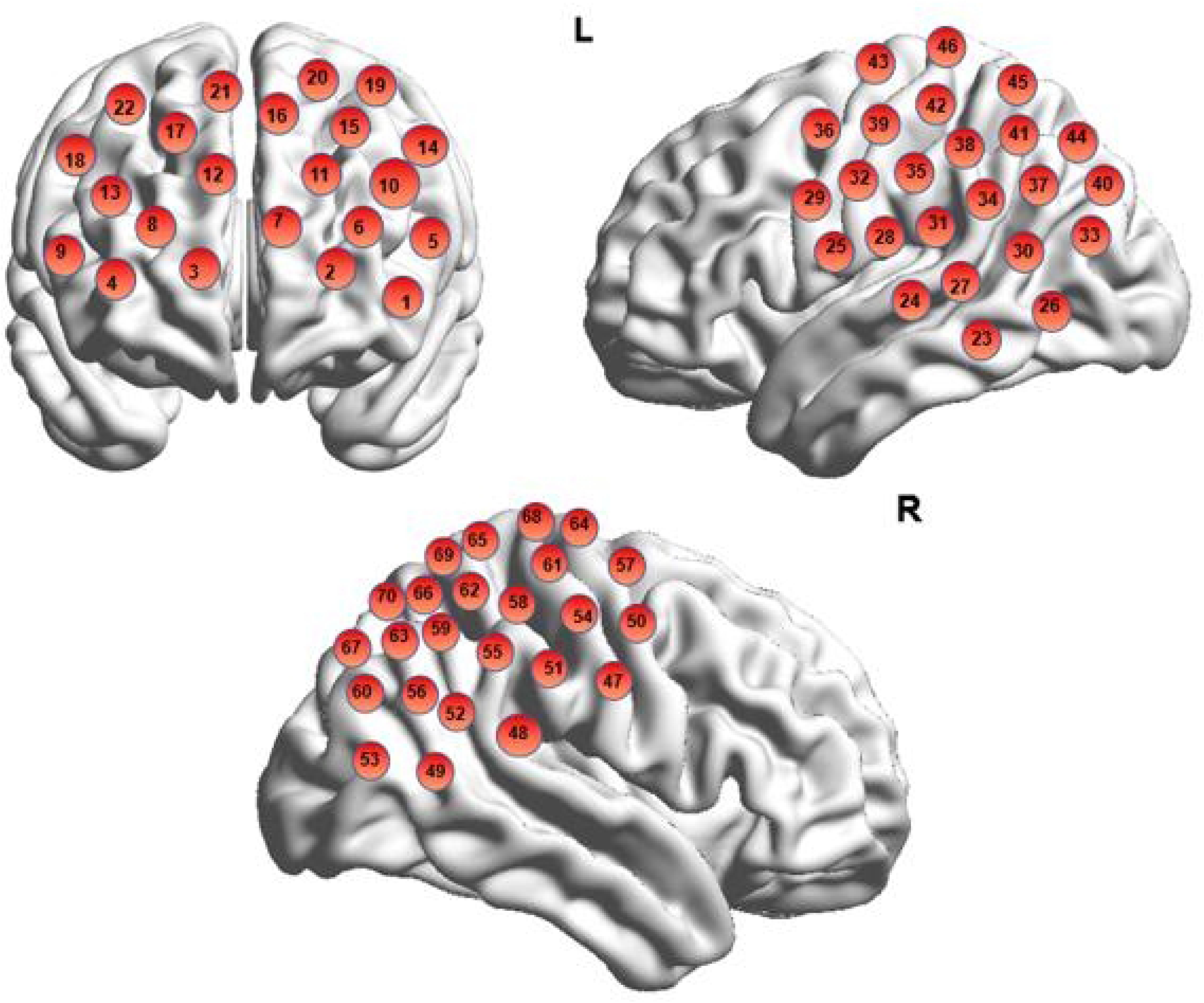
Schematic diagram of brain channel localization.

### fNIRS Preprocessing

Existing studies have shown that oxyhemoglobin (Hbo) is more sensitive to task-related stimuli^[38]^, so this study used MATLAB to observe only the changes in Hbo. To reduce task-irrelevant global signal interference, the HbO signal was preprocessed using the Principal Component Spatial Filtering algorithm (PCA)^[39]^, in order to eliminate the effect of global components. After that, the brain oxyhemoglobin signal was de-artifacted, including the removal of motion artifacts and instrumental noise, followed by spatial mapping of brain region activities using a source-space based distribution pattern method. After de-artifactualization of the acquired signals, the changes in blood oxygen levels in different brain regions under different decision-making conditions will be extracted. In order to ensure the quality of the data, the brain data were preprocessed using statistical tests to exclude abnormal data due to motion or equipment malfunctions.

### Statistical Analysis

This study used IBM SPSS 23.0 to analyze the data. First, demographic information and questionnaire data were analyzed by one-way analysis of variance (ANOVA) with group as a factor to examine differences between groups. Next, a 4 (groups: athlete stress group; general college student stress group; athlete sedentary group; general college student sedentary group) × 2 (time: pre-test; post-test) repeated measures ANOVA was performed on the heart rate and blood pressure data to test for indicators of significant main and interaction effects. Next, the behavioral data were subjected to a 4 (groups: athlete stress group; general college student stress group; athlete sedentary group; general college student sedentary group) × 2 (time: pre-test; post-test) repeated-measures ANOVA with post-hoc tests for significant indicators and tests for interaction effects. Then, based on the behavioral results, a 2 (group: athletes; general college students) × 2 (condition: stress; meditation) × 2 (time: pre-test; post-test) multifactorial ANOVA was conducted on the fNIRS data, and the channels with interaction and main effects were further analyzed. Finally, Spearman’s correlation analysis was applied to examine the association between behavioral changes and changes in brain activation.

## Results

### Descriptive statistical results

In order to explore the differences between the different groups on age, level of perceived fairness, experience of sense of power, and three-dimensional mood indicators of depression, anxiety, and stress. A one-way ANOVA was performed on the pre-test data, and it was found that there were no significant differences in any of the indicators (see Table 1).

**Table 1.**
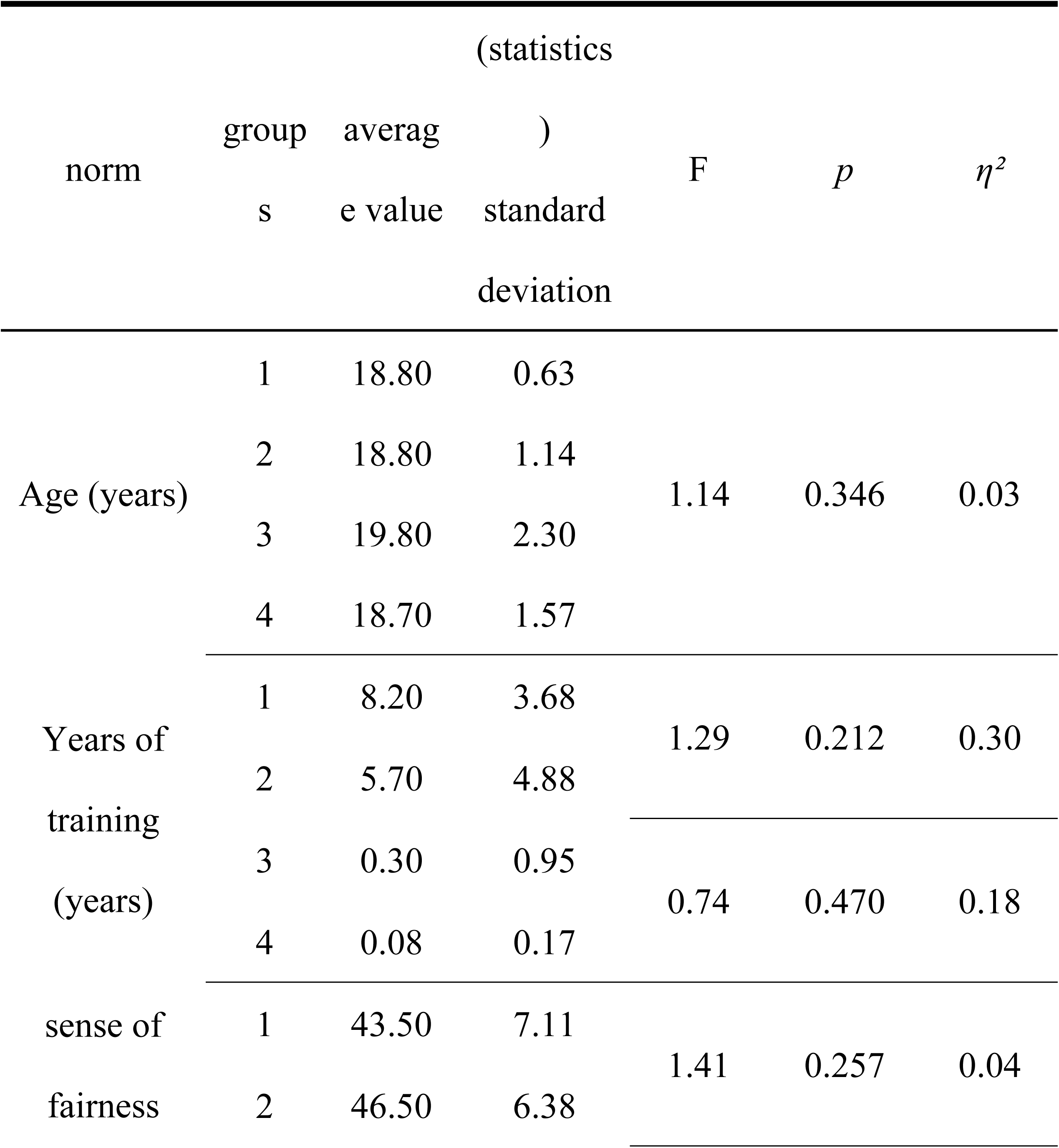
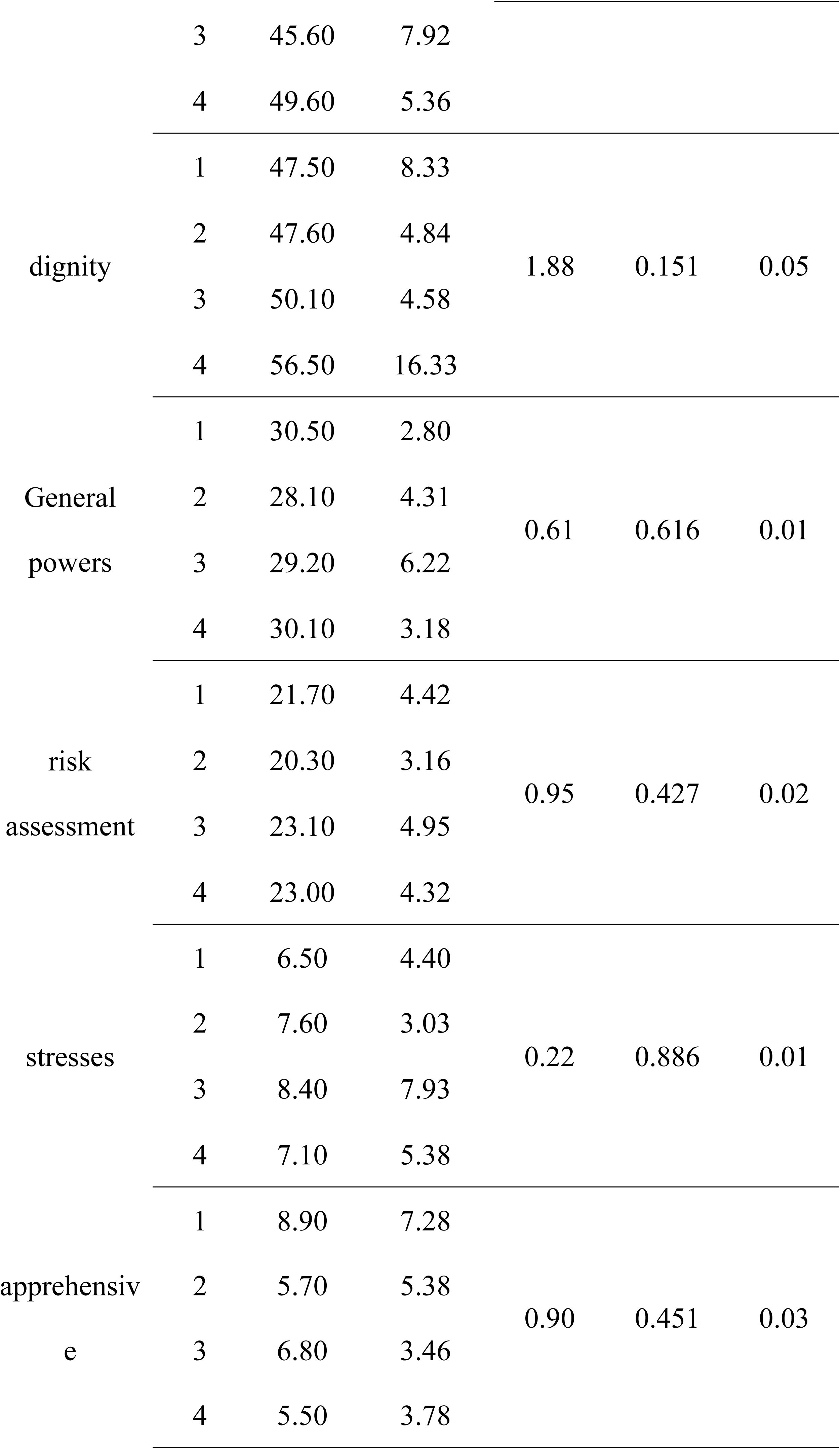
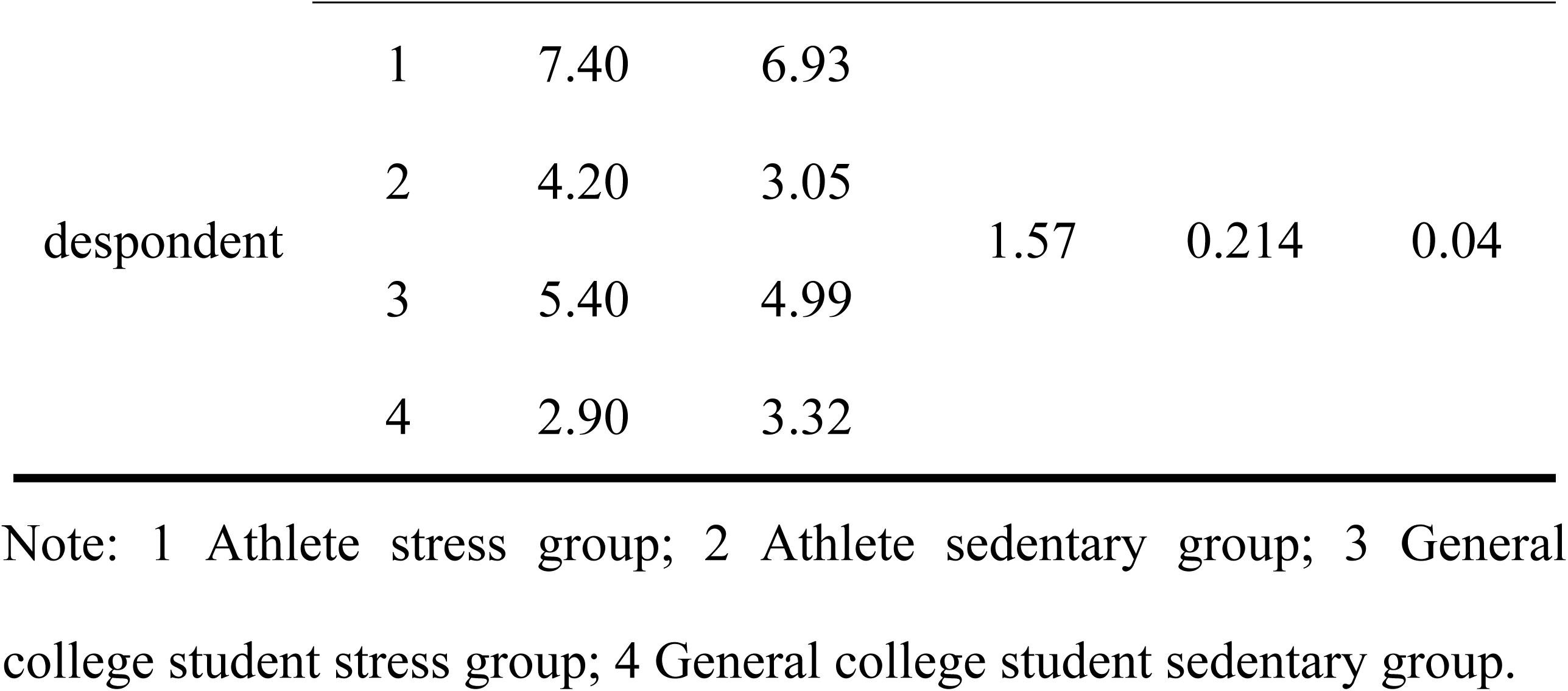
Results of descriptive statistics.

### Effect of acute stress on blood pressure and heart rate

In order to understand the effect of acute stress on blood pressure and heart rate, a 4 (groups: athlete stress group; general college student stress group; athlete sedentary group; general college student sedentary group) × 2 (time: pre-test; post-test) repeated measures ANOVA was performed. The results found that: in terms of blood pressure, there was no significant difference between the four groups in terms of systolic and diastolic blood pressure in the pre-test phase; in the post-test phase, there was a significant difference in terms of systolic and diastolic blood pressure between the athlete stress group and the athlete sedentary group (*p_intake_*= 0.003, *p_diastolic_*= 0.021), and there was a significant increase in systolic and diastolic blood pressure in the athletes after the stress (*p_intake_*= 0.010; *p_intraction_*= 0.058), while the systolic blood pressure of regular college students approached significant levels (*p* = 0.088) and diastolic blood pressure did not change significantly before and after stress. In terms of heart rate, no significant differences were found in either between-group comparisons or pre- and post-stress comparisons. The significant changes in blood pressure indicate that the effects of acute stress are effective in both athletes and regular college students.

### Effects of Acute Stress on Unfair Decision Making Behavior of Athletes

In order to investigate the effect of acute stress on unfair decision-making behavior, a 4 (groups: athlete stress group; general college student stress group; athlete sedentary group; general college student sedentary group)× 2 (time: pre-test; post-test) repeated measures ANOVA was conducted on the C1 extremely unfair scenario, the C2 relatively unfair scenario, and the C3 absolutely fair scenario, respectively (see Appendix-Table 1). The results found significant time main effects for the three programs _(*p(C1-refusal) <0*_.001, _*p(C2-refusal) <0*_.001, _*pC2-takeover*_=0.005, _*p(C3-takeover) <0*_.001), and group main effects (_*pC2-refusal*_= 0.002, _*pC2-takeover*_= 0.033) and interaction effects for the relatively inequitable program were significant _(*p(C2-reject)*_ <0.001, _*pC2-connect*_= 0.001).

In order to compare the differences between athletes and average college students in their decision-making on the sense of unfairness, a post hoc test on the pre-test data found that there were no significant differences between athletes and average college students in the rejection and acceptance efficiencies of the three scenarios ^(*p*^*C1-Reject*^=0.397,*p*^*C1-Receive*^=0.751,*p*^*C2-Reject*^=0.054,*p*^*C2-Receive*^=0.248,and^ *p_C3-Refuse_*=0.417, *p_C3-Receive_*=0.138), suggesting that athletes and regular college students are consistent in their decision-making about feelings of unfairness.

In order to investigate the effect of acute stress on athletes’ decision-making on sense of unfairness, a simple effects analysis was conducted on the relative unfairness scenarios with interaction effects. The results found that the acceptance efficiency of the posttest in the athlete stress group was significantly higher than that of the athlete meditation group (*p*<0.001) and the general college student stress group (*p*<0.01), and significantly higher than that of the pretest (*p*<0.001) (see Figure 4). On the rejection efficiency, the difference between the pretest and post-test in the athletes’ stress group was not significant (*p>* 0.05). The above results indicate that acute stress can significantly affect athletes’ decision-making on the sense of relative unfairness, and athletes will be more inclined to make the decision of accepting the relative unfairness program after stress compared with ordinary college students.

**Figure 4.**
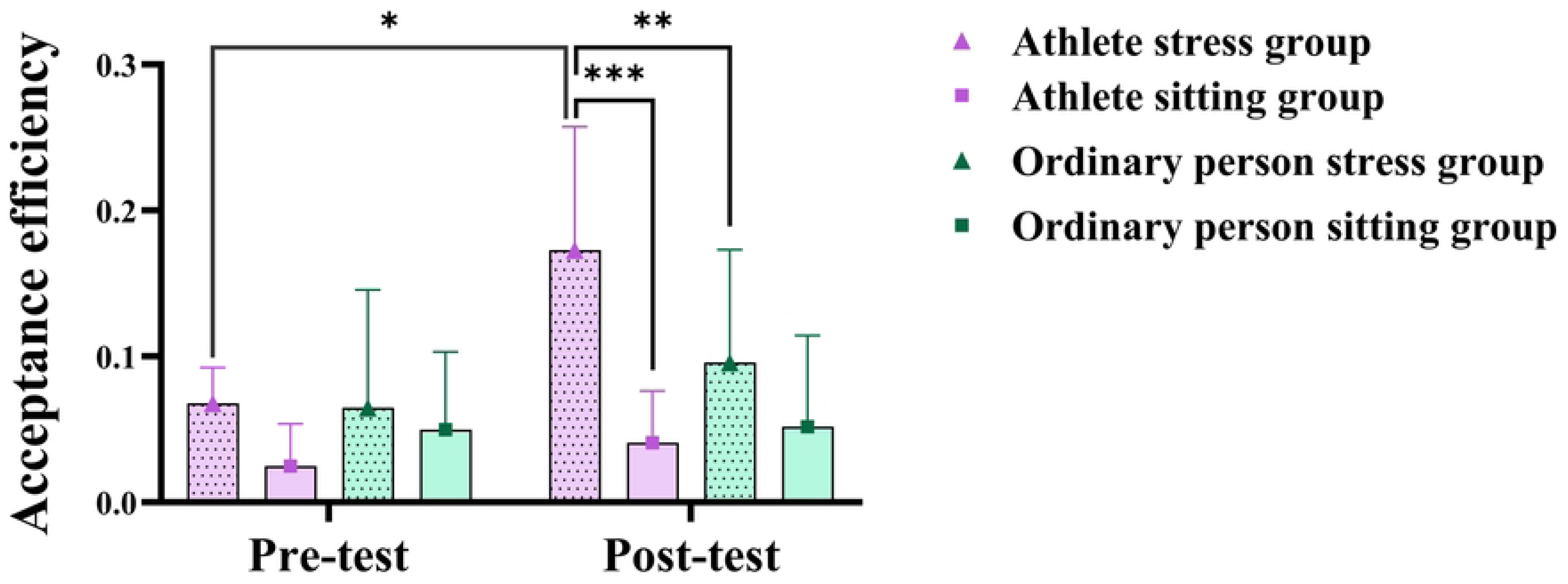
Three-way ANOVA results of acute stress on relatively unfair decision-making behavior.

### Effects of Acute Stress on Brain Activation in Athletes Undergoing Relatively Unfair Programs

To investigate the brain mechanisms by which acute stress affects athletes’ acceptance of relatively unfair programs, a three-factor mixed design of 2 (group: athletes vs. regular college students) × 2 (condition: stress vs. sedentary) × 2 (time: pre-test vs. post-test) was used, and ANOVA showed that on CH15 (located in the Frontal Eye Fields, frontal eye area, Brodmann 8,see Figure 5A), a significant group x condition x time interaction effect, and further post hoc tests revealed that activation on the post-test was significantly lower in the athlete stress group than on the pretest (*p*=0.009) and significantly lower than on the post-test in the general college student stress group (*p*=0.013); whereas activation on the post-test in the general college student sedentary group was significantly lower than on the pretest (*p*=0.017) (see Figure 5A).

**Figure 5.**
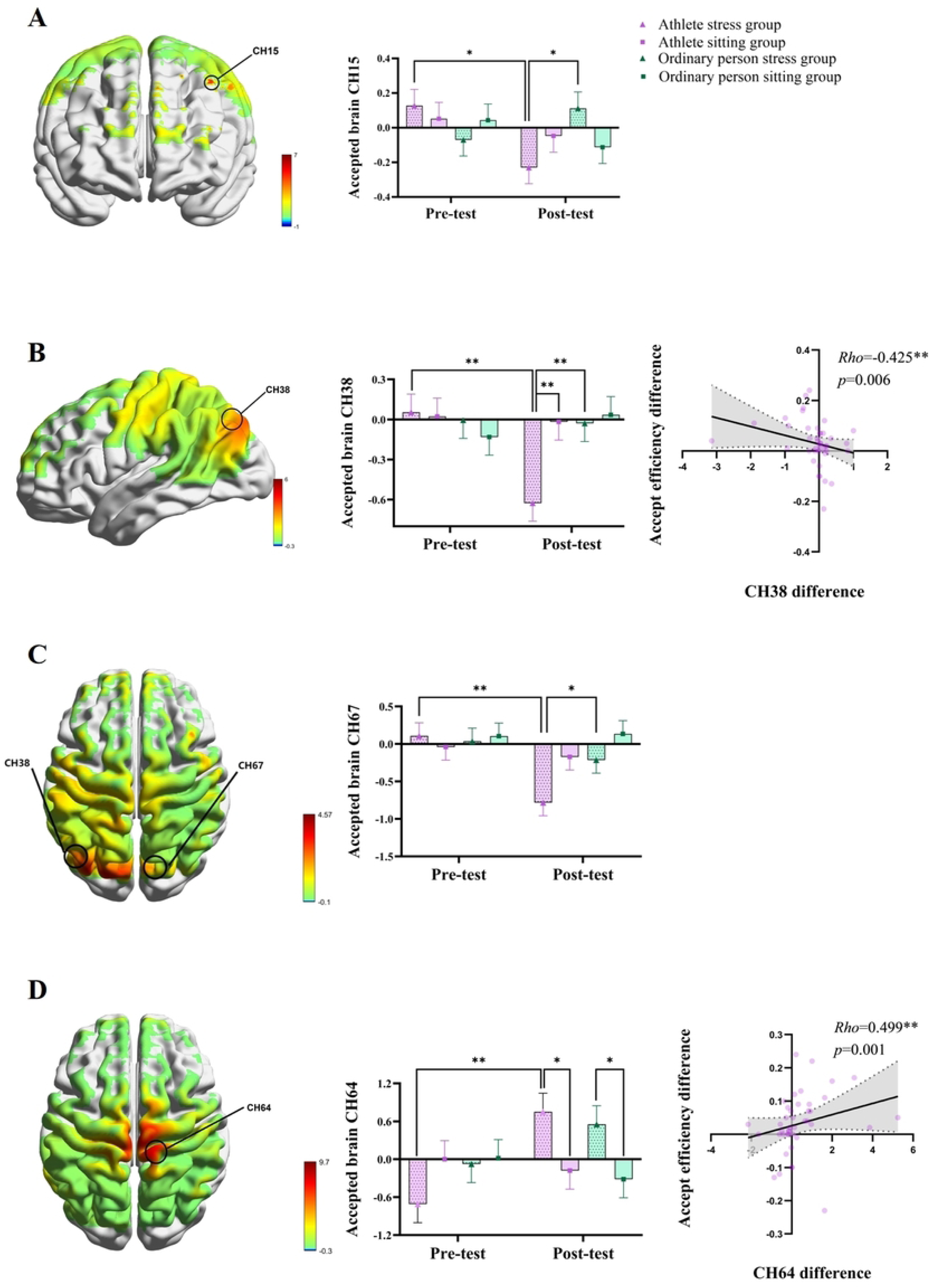
A. Mapping of group × condition × time interaction effect.

On CH38 (located in the Supramarginal gyrus, the supramarginal gyrus, Brodmann 40,see Figure5B), the group × time, and condition × time interaction effects were significant, and further post-hoc tests revealed that activation was significantly lower in the athlete stress group post-test than in the pre-test (*p*=0.001), and was significantly lower than in the athletic sedentary group post-test (*p*=0.002) compared to the general college student stress group posttest (*p=0*.003). On CH67 (located in Somatosensory Association Cortex, Somatosensory Joint Cortex, Brodmann 7,see Figure5C), there was a significant group × time interaction effect, and further post-hoc tests revealed that activation was significantly lower in the athlete stress group post-test than in the pre-test (*p*=0.001) and significantly lower than in the general college student stress group post-test (*p*=0.025). On CH64 (located in Primary Motor Cortex, Primary Motor Cortex M1, Brodmann 4,see Figure5D), the condition × time interaction effect was significant, and further post-hoc tests revealed that activation in the athlete stress group post-test was significantly higher than in the pre-test (*p*=0.001) and significantly higher than in the exercise sedentary group post-test (*p*=0.028); whereas, activation in the general college student stress group post-test was significantly higher than in the general college student sedentary group post-test activation was significantly higher than the posttest of the general college students’ sedentary group (*p*=0.040). The above results indicated that acute stress caused a significant decrease in CH15, CH38 and CH67 channel activation and a significant increase in CH64 channel activation in athletes undergoing relatively unfair decision-making.

To further explore the covariate relationship between decision-making behavior and brain function, Spearman’s rank correlation test was used to examine the difference between pre and post-intervention behavioral and brain activation differentials. The results showed that there was a moderate negative correlation between the difference in acceptance efficiency of athletes before and after acute stress and the difference in CH38 brain activation (Rho=-0.425, *p*=0.006) and a moderate positive correlation with the difference in CH64 brain activation (Rho=0.499, *p*=0.001). The above results suggest that behavioral changes in athletes undergoing relatively unfair protocols after acute stress are closely related to reduced activation of CH38 and elevated activation of CH64.

F-values on the brain & specific change in CH15; B. Mapping of group × time interaction effect F-values on the brain & specific change in CH38 & correlation of change in CH38 with change in receptive efficiency; C. Mapping of group × time interaction effect F-values on the brain & specific change in CH67; D. condition × time interaction effect F-value mapping on the brain & specific change in CH64 & correlation of change in CH64 with change in receptive efficiency

## Discussion

The present study employed a randomized controlled experimental design using near-infrared spectroscopic imaging (fNIRS) and an ultimatum gaming (UG) task with the Maastricht Stress Test (MAST) to induce acute stress, aiming to investigate the behavioral performance of athletes and ordinary college students and their blood oxygen signal changes in the prefrontal, right and left temporoparietal joint regions when they completed the unfairness sense decision-making task after acute stress. The results of the study revealed that acute stress can significantly affect athletes’ sense of unfairness decision-making, and athletes were more likely to accept a relatively unfair solution after stress compared with ordinary college students; this behavioral performance was associated with reduced activation of CH38 (supramarginal gyrus, Brodmann 40) and elevated activation of CH64 (primary motor cortex M1, Brodmann 4).

This study found no significant difference in unfairness - related decision - making between athletes and non - athlete college students. We speculate that such decision - making may be influenced by social norms and the environment. The athletes recruited were university - based, exposed to the same campus culture and education as non - athletes, leading to similar basic perceptions and expectations of fairness. However, after acute stress, athletes showed a stronger acceptance of relatively unfair proposals than non - athletes. We propose that athletes, often under intense training and competitive pressure, develop unique decision - making strategies to cope with stress. For instance, successful table tennis players often use avoidance strategies during matches.Studies on golfers also indicate that task - oriented approaches help achieve goals, enhance psychological adjustment, and boost decision - making skills^[40, 41]^. These studies show that long - term training and competition enable athletes to quickly adjust decision - making strategies under stress, adapt to the environment, and make more self - favorable choices. Our findings on athletes’ unfairness - related decision - making align with prior studies. Under stress, athletes rapidly adjusted their strategies to accept relatively unfair proposals. This indicates they have higher post - stress tolerance for unfair events. They turn “compromise” into a learned adaptive behavior. This isn’t due to exhausted cognitive resources but is a proactive strategy adjustment. It’s similar to tactical in - match compromises, possibly to achieve short - term goals or maintain performance. In contrast, non - athletes can’t adjust decision - making as effectively under stress. These results are highly significant for understanding psychological and behavioral responses to stress across groups and offer valuable insights into the mechanisms of stress - induced decision - making..

Knoch et al. (2008) used transcranial direct current stimulation (tDCS) to inhibit the right dorsolateral prefrontal cortex and found that participants accepted unfair behavior more often, showing the direct impact of specific brain - area activity changes on decision - making regarding unfairness^[42]^. In our study, after stress, athletes showed reduced activation in the supramarginal gyrus and increased activation in the primary motor cortex. These neural changes correlated with athletes’ increased acceptance of relatively unfair proposals post - stress. The supramarginal gyrus, key for social cognition, emotion regulation, and decision - making, saw reduced activation, suggesting athletes had lowered sensitivity to unfairness or changed emotion regulation under stress. This shift reduced over - emphasis on fairness, focusing more on behavioral feasibility. The primary motor cortex’s increased activation might relate to athletes’ physiological responses and behavioral preparation under stress, enabling faster information integration and decision - making. These neural changes indicate that athletes’ decision - making advantage under stress stems from long - term training and competition - induced biological adaptation. Our findings align with Knoch’s study, highlighting the importance of specific brain - area activity in decision - making, especially in unfair situations, where reduced activation in certain areas may lower sensitivity to unfairness.

While this study highlights the acute stress impact on athletes’ unfairness - related decision - making and links it to specific neural activity changes, some limitations remain. For instance, it only examines the immediate effects of acute stress, not tracking behavioral and brain - function dynamics post - stress recovery. Future studies could use multi - time - point designs (e.g., 0, 30, and 60 minutes post - stress) to explore the timeliness of stress effects and recovery mechanisms. Also, this study doesn’t differentiate between sport types (team vs. individual). Given potential differences in stress - adaptation mechanisms among athletes of various disciplines, future research can increase sample sizes and stratify by sport type and training duration to investigate group heterogeneity in stress responses.

## Conclusion

The study found that athletes tend to accept relatively unfair solutions after acute stress and that this behavioural performance is associated with reduced activation in the supramarginal gyrus and elevated activation in the primary motor cortex. The present study provides new perspectives for understanding the psychological and neural mechanisms of athletes in stressful situations, and also provides important theoretical references for athletes’ psychological training and competition strategy development.

## Data availability statement

All datasets generated for this study have been included in the manuscript and/or supplementary files.

## Source of funds

This study was funded by the Shanghai Innovative Training Programme for Undergraduates (Project No. 202510269085S).

